# Pathway and directional specificity of Hebbian plasticity induction in the cortical visual motion processing network

**DOI:** 10.1101/2022.05.15.491882

**Authors:** Michele Bevilacqua, Krystel R. Huxlin, Friedhelm C. Hummel, Estelle Raffin

**Affiliations:** Defitech Chair in Clinical Neuroengineering, Center for Neuroprosthetics and Brain Mind Institute, EPFL, Geneva, Switzerland; Defitech Chair in Clinical Neuroengineering, Center for Neuroprosthetics and Brain Mind Institute, Clinique Romande de Readaptation (CRR), EPFL Valais, Sion, Switzerland; The Flaum Eye Institute and Center for Visual Science, University of Rochester, Rochester, NY, USA; Clinical Neuroscience, University of Geneva Medical School, Geneva, Switzerland

**Keywords:** Visual motion processing, TMS-EEG, Motion discrimination, Cortico-cortical Paired Associative Stimulation, Granger Causality

## Abstract

Cortico-cortical paired associative stimulation (ccPAS), which repeatedly pairs single pulse TMS over two distant brain regions with a specific time interval, is thought to modulate synaptic plasticity. Applied to the motion cortical pathway, ccPAS has been shown to improve motion discrimination when specifically targeting backward projections, stimulating the medio-temporal area (MT) followed by the primary visual cortex (V1). However, there is no direct neuroimaging evidence of the spatial selectivity of the ccPAS effects (i.e., pathway or direction specificity) or detailing the exact nature of the ccPAS effects (i.e., the oscillatory signature, timing…). In this study, we applied ccPAS along the motion discrimination pathway, in the top-down direction (MT-to-V1: “Backward ccPAS”) and in the bottom-up direction (V1-to-MT: “Forward ccPAS”) in sixteen healthy volunteers and compared changes in visual network activity in response to single pulse TMS over V1 and MT using spectral granger causality (sGC). The sGC results showed common increases in direct V1-to-MT and V1-to-IPS bottom-up inputs in the high Beta/low Gamma band (25-40 Hz) for both ccPAS, probably reflecting task exposure. However, a clear distinction in information transfer occurred in the re-entrant MT-to-V1 signals, which were only modulated by Backward ccPAS. This difference was predictive of the behavioural improvements at the motion discrimination task. Our results support the view of the possibility to specifically enhance re-entrant Alpha oscillatory signals from MT-to-V1 to promote motion discrimination performance through Backward ccPAS. These findings contribute to better understanding visual processing in healthy subjects and how it can be modulated to pave the way to clinical translation in vision handicapped patients. The changes in re-entrant MT-to-V1 inputs could help to provide single-subject prediction scenarios in patients suffering from a visual system stroke, in whom visual recovery might partly rely on the top-down inputs to the spared V1 neurons.

## Introduction

Visual processing represents a massive computational task for the brain, requiring highly organized and efficient neural systems to achieve accurate perception. In primates, approximately 55% of the cortex is somehow dedicated to visual processing (compared to 11% for somatosensory processing or 3% for auditory processing) (Felleman et al., 1991). The different visual pathways have to receive, relay, and ultimately process visual information. These structures include the eye, retina, optic nerves, chiasm, tracts, lateral geniculate nucleus (LGN) of the thalamus, radiations, the primary visual cortex (or striate cortex), and the secondary visual cortex (extrastriate cortex). Information processing relies not only on forward connections to higher visual areas, but also on reciprocal connections that transfer information in the opposite direction. These backward connections, which are much more numerous than the forward inputs coming from the LGN (Catani et al., 2003), are thought to adjust, regulate and improve the processing of incoming stimuli (Rynolds et al., 2003; Antal et al., 2004; Bressler et al., 2008). Therefore, neural responses in V1 do not mirror retinal inputs precisely, but are modified by higher inputs to support a coherent perceptual interpretation.

One of the most studied cortical visual processing pathways is the one specialized in decoding motion stimuli (Mikami et al., 1986). The very early stage involves the primary visual cortex (V1). Early single unit recordings in macaque and cats showed that a subset of cells in V1 are highly direction selective (Hubel et al., 1968; Gizzi et al., 1990; Maunsell et al., 1983). These direction sensitive cells then project onto the medial temporal area (MT), where all neurons appear to be directionally selective (Maunsell et al., 1983; Albright et al., 1984; Mikami et al., 1986). More precisely MT neurons are capable of coding the direction and speed of the image motion (Felleman et al., 1984; Lagae, et al., 1993; Perrone et al., 2001; Rodman et al., 1987). Lesion studies on cortical blindness patients or macaques (Pasternak et al., 1993; Das et al., 2014) as well as transcranial magnetic stimulation (TMS) studies using virtual lesion approaches on humans (Ellison et al., 2003, Cowey et al., 2001) further provided evidence supporting the role of both areas in motion discrimination. For instance, Marcar *et al*. (1992) trained nine macaques at a random dot motion task. The subsequent surgical removal of the MT complex in two monkeys induced a loss of their ability to discriminate coherently moving dots. The majority of human studies have concentrated on the capacity of MT cells to register motion in the fronto-parallel plane and on their directional properties and speed preferences (Maunsell et al., 1983; Rodman et al., 1987; Priebe et al., 2003; Liu et al., 2005). In this regard, Ruzzoli *et al*. (2010) studied the effect of a TMS perturbation on the shape of the psychometric function in a visual motion discrimination task. The results showed a decrement in motion discrimination performance when TMS was applied over MT concomitantly to the motion stimulus. Furthermore, the authors found that TMS specifically affected the perceptual ability to discriminate motion direction rather than higher cognitive processes such as perceptual consciousness, decision-making, or response selection and execution.

The interaction and synchronization between visual areas have been thoroughly investigated in the past years in different models. One of the first computational model that simulates the dynamic integration in the visual system and the pivotal role of backward connections between the visual areas in motion perception processing was proposed by Tononi *et al*. (1992). The model reported that re-entrant connections in the visual cortex are mostly integrative, i.e., they facilitate the coordination of neuronal firing in anatomically and functionally segregated cortical areas. Thus, neural responses related to the detection of motion coherence may be depending on the re-entrant projections from area MT to area V1, that have been found to be mostly excitatory (Pan et al., 2021). The timing of the backward inputs from MT to V1 has been studied in humans using visual masking stimuli (e.g., Breitmeyer et al., 2006), visually-evoked potentials (e.g., Wibral et al., 2009) or by TMS pertubation (e.g., Pascual-Leone et al., 2001, Silvanto et al., 2005, Laycock et al., 2015). TMS studies targeting MT have consistently shown two periods of disruption suggestive of forward/backward models (e.g., Laycock et al., 2007). Furthermore, paired-pulse TMS protocols can estimate the time delay for information transfer along the pathway. For instance, Pascual-Leone *et al*. (2001) showed significant impairments of coherent perceptual interpretation of motion when a MT pulse was given 10 to 30 ms before a V1 pulse (Pascual-Leone and Walsh, 2001). Importantly, this processing channel possesses characteristic oscillatory activity that is linked to the transfer of specific stimulus features or endogenous variables within the pathway. Backward projections have been found to be mediated by low frequency Alpha/Beta oscillations while forward connections are thought to be supported by high frequency Gamma oscillations as shown in monkeys (van Kerkoerle et al., 2014, Bastos et al., 2015) and in humans (Michalareas et al., 2016).

An ultimate approach to non-invasively infer on the role of backward projections for motion discrimination is to causally manipulate the synaptic strength of the top-down MT-to-V1 connection. This is possible through the use of cortico-cortical Paired Associative Stimulation (ccPAS) (Chiappini et al. 2018). This approach relies on the Hebbian Learning theory, which states that the pairing of subthreshold synaptic stimulation and action potential trains can induce LTP (Long-Term Potentiation) (Magee et al. 1997). The first human applications performed by Stefan *et al*. (2000), showed that low-frequency TMS over the primary motor cortex following a peripheral stimulation of the median nerve induces plastic changes in the human motor system when using timings relevant for spike-timing-dependent plasticity (STPD) (Caporale et al., 2008). This concept has been applied to cortico-cortical connections, for instance between frontal and parietal areas (Koch et al., 2018), or between the cerebellum and M1 (Spampinato et al., 2017). ccPAS has also been applied to the V1-MT pathway (Romei et al., 2016). These authors compared various versions of ccPAS between V1 and MT on healthy subjects and found that only ccPAS targeting the re-entrant connection from MT to V1 with an optimal time delay of 20ms was effective in boosting motion discrimination capacities up to 90 minutes.

In the present study, we applied the most efficient ccPAS configurations, based on literature, to the V1-MT pathway to compare the effects of enhancing forward versus backward projections on motion discrimination and visual network activity. We used EEG, and spectral Granger Causality (sGC) based network analyses to investigate pathway specificity and directional specificity of ccPAS, as well as the spectral content of the induced changes. The combination of sGC and single pulse TMS over V1 and MT allowed us to distinguish output connectivity from re-entrant connectivity (Keil et al., 2009). In line with a previous study (Romei et al., 2016), we hypothesized that enhancing backward projections would induce larger behavioural improvements and would be associated with a specific increase in connectivity restricted to the re-entrant backward MT to V1 inputs, especially in the Alpha band. Conversely, we hypothesized that enhancing forward inputs would induce non-specific changes in bottom-up direct output connections, not necessarily relevant for motion perception.

## Methods

### General procedure and participants

16 healthy subjects (9 F, 7M, mean age: 27±4 yo±SD) participated in the study. The double-blinded and cross-over design of the study, implied two similar sessions of 3 hours each, only differing by the type of ccPAS intervention applied to the participants. The two sessions were performed at least one month apart and the order was randomized and counter-balanced between participants. Each session comprised a familiarization phase including 3 blocks of 20 trials each, to ensure that the subjects understood the visual discrimination task and reached stable performance. After EEG cap preparation, TMS sites and intensities were defined (see below for more details). Task performances and EEG responses to single pulses over V1 and MT were measured at baseline, followed by one of the two ccPAS interventions. Ten minutes after the end of the ccPAS, task performances and single pulse TMS-EEG over V1 and MT were measured again. During the whole experiment, the participants sat on a chair, with the head leaning on the chinrest, in front of a computer screen, centred 47 cm far from the eyes. Every subject was asked to fill in a short questionnaire related to eventual issues and inconveniences caused by the single pulse TMS or ccPAS intervention at the end of both sessions. All participants provided informed written consent prior the experiment and none of them met the MRI or TMS exclusion criteria (Rossi et al., 2021). This study was approved by the local Swiss Ethics Committee (2017-01761) and performed in accordance with the Declaration of Helsinki.

### Single pulse TMS

For single pulse TMS over V1 and MT, biphasic TMS pulses inducing an antero-posterior followed by postero-anterior current in the brain (AP-PA) were sent through a MC-B65-HO butterfly coil (MagVenture A/S, Denmark) plugged in a MagPro XP TMS stimulator (MagVenture A/S, Denmark). TMS was delivered over V1 and the right MT using the Localite neuronavigation software (Localite GmbH, Germany). To precisely target the individual V1 and MT areas, we used a standard fMRI MT localizer task performed prior the TMS-EEG session (Figure 1B). During the functional localizer, the screen displayed radially moving dots alternating with stationary dots (see e.g., (Sack et al., 2007)). A block design alternated six 15 s blocks of radial motion with six blocks featuring stationary white dots in a circular region on a black background. This region subtended 25° visual angle, with 0.5 dots per square degree. Each dot was 0.36° diameter. In the motion condition the dots repeatedly moved radially inward for 2.5 s and outward for 2.5 s, with 100% coherence, at 20°/s measured at 15° from the center. Participants were passively looking at the screen and were asked to focus on a fixation point located in the middle of the screen. The resulting activation map and the individual T1 image were entered into the neuronavigation software to define the coil positions as seen in Figure 1A. The mean coil positions for V1 were −16±9, −86±8, −5±22 and for MT were 66±9, −55±9, −7±17 (coordinates x, y z, MNI space). The coil was held tangentially to the scalp with the handle pointing upwards and laterally at 45° angle to the sagittal plane.

**Fig. 1:**
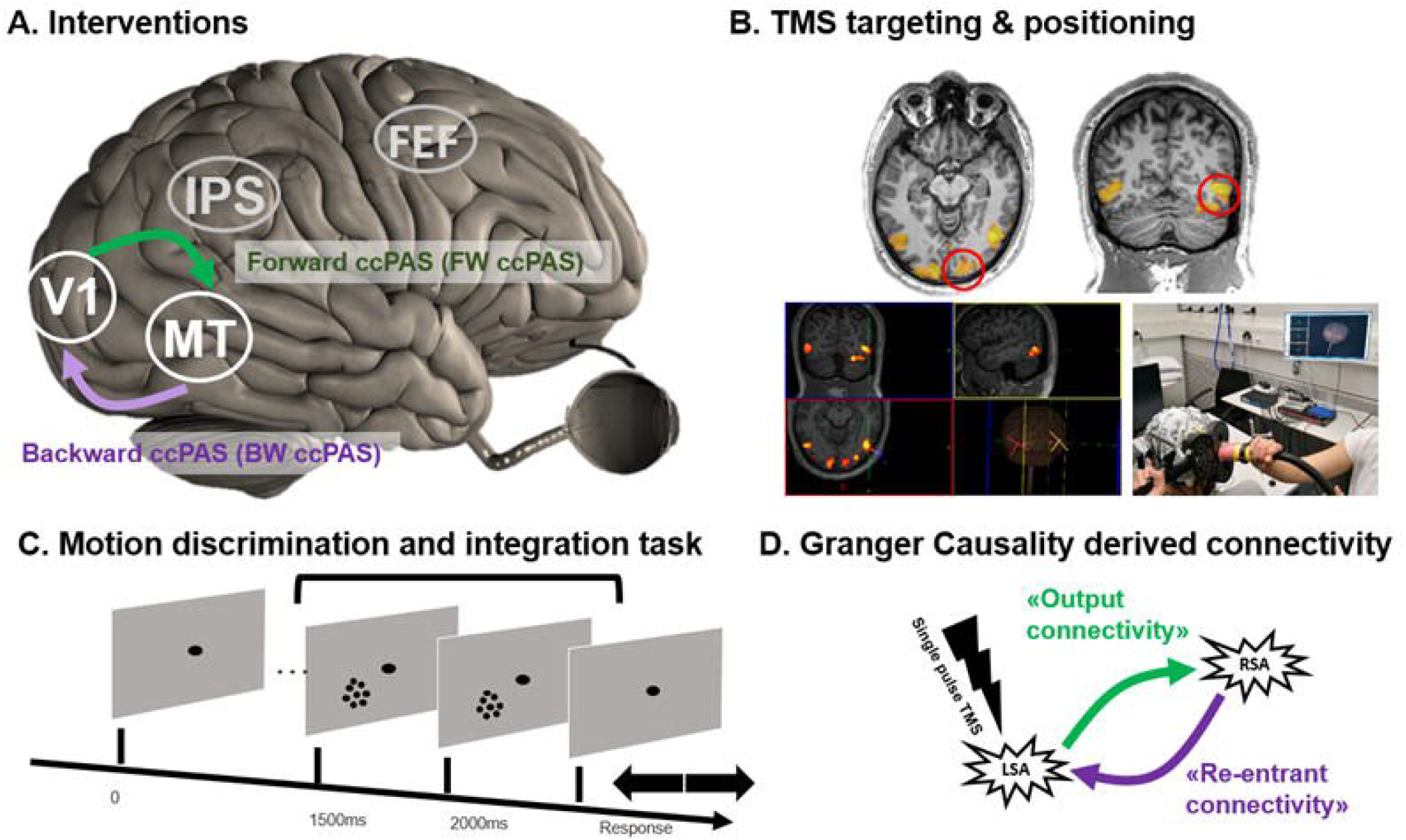
A: Illustration of the Backward (BW) and Forward (FW) ccPAS interventions as well as the four regions of interest; **B**: Example of an individual functional localizer, online neuronavigation and coil positioning; **C:** Illustration of the motion direction discrimination and integration task used before and after the ccPAS interventions; **D**: Illustration of the local and remote source activity (LSA and RSA) as well as the dissociation between output and re-entrant Granger Causality derived connectivity estimates between V1 and MT.

To determine stimulation intensities, we evaluated the phosphene threshold (Gerwig et al., 2003) on both V1 and MT. If the participants reported phosphenes (9/16 for V1 and 5/16 for MT), we set the stimulation intensity at 90% of the PT. If no phosphenes could be evoked, we used 65 % MSO in all participants to maximize signal-to-noise ratio and comfort during the exam. Because of discomfort related to the fact that MT is located close to the ear we decrease the intensity to 60% MSO. The mean stimulation intensity for V1 was 66±6 % MSO and for MT 61±4 % MSO. On each area, 90 TMS pulses were performed with an inter-pulse interval of 4±1s.

We used active noise cancellation intra-auricular earphones (Bose QC 20, USA) to mask the TMS click susceptible to evoked auditory responses on the ongoing EEG activity. The sound level was adjusted for each subject, so that the TMS click delivered became barely audible without any discomfort for the participant. A thin layer of soft plastic was placed on the coil surface to dampen both sensory and auditory feedbacks to the subject.

### ccPAS interventions

ccPAS was delivered via two independent TMS stimulators externally triggered with Signal (Digitimer, Cambridge Electronic Design, Cambridge, UK). V1 was stimulated with a MagVenture MagPro XP stimulator (MagVenture A/S, Denmark) connected to a MC-B65-HO coil and the right MT area was stimulated with a MagVenture MagPro X100 stimulator connected to a B35 coil using the same coil positions as for single TMS (see Figure 1B). The same intensities were used for MT. For V1, since we used a different coil, we recalibrated the stimulation intensity to match the single pulse TMS condition. 90 pairs of biphasic stimuli were continuously delivered at a rate of 0.1 Hz for 15 min (same as Romei et al., 2016). Online navigation of both coils was used throughout the intervention. The coil handle for MT was held tangentially to the scalp and pointed downwards at an angle of 120°±5 clockwise. Because our goal was to explore the direction-specificity effects of ccPAS in the motion processing network, we compared two types of ccPAS, differing by the order of the two pulses. In the MT-V1 Backward ccPAS condition *(BW ccPAS*), the first TMS pulse was applied to MT followed by another pulse to V1. In the V1-MT Forward ccPAS condition *(FW ccPAS)*, the TMS pulses order was reversed; the first pulse was administered to V1 and the second to MT (*Fig. 1D*). The inter-stimulus interval (ISI) was set at 20ms for both ccPAS conditions, because it corresponds to the time delays of MT-V1 back projections (Pascual-Leone et al. 2001; Silvanto et al. 2005). This timing is critical to create sequential presynaptic and postsynaptic activity in the network, and to finally generate the occurrence of STDP (Caporale et al. 2008; Jackson et al. 2006).

### Behavioural task

We used the same 2-alternative, forced-choice, left-right, global direction discrimination and integration task (150 trials in total), as previously described (Huxlin et al., 2009; Martin et al., 2010; Salamanca-Giron et al., 2021, Raffin et al., 2021). The stimuli consisted of a group of black dots moving globally left-or rightwards at a density of 2.6 dots/° and in a 5° diameter circular aperture centred at cartesian coordinates [-5°, 5°] (i.e., the bottom left quadrant of the visual field, relative to central fixation, see Figure 1D).

The range of dot directions was varied between 0° (total coherence) and 360° (complete random motion) in steps of 40°. The degree of difficulty or direction range was increased with improving task performance by increasing the range of dot directions within the stimulus. A 3:1 staircase design was implemented. For every 3 consecutive correct trials, direction range increased by 40°, while for every incorrect response, it decreased by 40°. The black dots forming the stimulus were 0.06° in diameter and moved at a speed of 10°/s over a time lapse of 250ms for a stimulus lifespan of 500ms. At every stimulus onset, an auditory beep was played for the subject. Self-confidence was rated (low/medium/high) after each trial. After each trial, auditory feedback indicated whether the response was correct or incorrect.

The task was implemented in Matlab 2019b (Version 1.8) (The MathWorks Inc., USA) coupled with an EyeLink 1000 Plus Eye Tracking System (SR Research Ltd., Canada) sampling at a frequency of 1000 Hz to control gaze and pupils’ movements in real time. the task was projected onto a mid-grey background LCD projector (1024 × 768 Hz, 144 Hz) and participants used the left-right arrows of the keyboard with their right hand and the “a, s and d” (for low/medium/high confidence), for the trial-by-trial confidence.

### EEG recordings

EEG was recorded using a 64 channels TMS compatible system (BrainAmp DC amplifiers and BrainCap EEG cap, Brain Products GmbH, Germany) with ground electrode in Fpz, reference electrode in Cz and Iz electrode added to the international standard 10-20 layout. Electrode impedances were adjusted and kept under 5 kOhms using conduction gel. The impedance levels were checked throughout the experiment and corrected if needed during breaks between the recordings. The signal was recorded using DC mode, filtered at 500 Hz anti-aliasing low-pass filter and digitalized at 5 kHz sampling frequency. Channel coordinates were individually assessed using the neuronavigation software at the end of the experiment.

### TMS-EEG Preprocessing

EEG analysis have been performed on the EEG recordings of the 90 single pulses sessions over V1 and MT. All the preprocessing have been computed on MATLAB, using the EEGLAB toolbox, the open source TMS-EEG Signal Analyser (TESA) plugin (Rogasch et al. 2017) and the plugin Brainstorm (Tadel et al., 2011).

Detection of bad channels was performed using the EEGLAB built-in function. Then, the raw EEG signal was epoched in a window of [-0.2, 0.8] s around the stimulation pulse onset, and demeaned. Afterwards, a window of [-5, 25]ms around the pulse onset was removed, in order to remove the TMS artefact and the missing data was interpolated using a cubic function by considering the data 5ms before and after the removed TMS artefact window. EEG data were then down-sampled from 5000Hz to 1000Hz, and bad epochs (the one presenting huge rubbing artefacts or undefined significant noise) were removed by visual inspection. On average, 13 epochs out of 90 were removed per recording. Interpolated data in the TMS artefact window were removed gain, in order to compute the first round of Independent Component Analysis (ICA), aiming at removing components of the pulse artefact. This first ICA is computed by first performing a PCA compression using the TESA built-in function, and by then performing a symmetric fastICA with hyperbolic tangent as contrast function. The artefact components were removed manually by visual inspection. On average, 7 (±3) components out of the 64 were removed. The EEG signal was re-interpolated in the removed window as done previously, and frequency filtered with a bandpass filter between 1 and 80Hz, with a 4th degree Butterworth notch filter between 48 and 52Hz to remove the power line noise. Data belonging to the artefact window were removed for the last time as done previously, and a second round of fastICA was in order to remove the other type of artefacts, as eye movement, blinking, acoustic artefacts and small head movement (Rogasch et al. 2017). On average, 4 (±1) components over 64 were removed. Finally, the EEG signal was spatially filtered using Common Average Reference (CAR) filter.

### TMS-EEG Local/Remote Source Activity (L/RSA)

Source reconstruction for each TEP was performed following the default procedure proposed in Brainstorm (version 23-Mar-2022) software (Tadel et al., 2011) together with OpenMEEG BEM plugins. First, the cortex and head meshes (15,000 and 10,000 vertices respectively) of each individual were generated using the automated MRI segmentation routine of FreeSurfer (Reuter et al., 2012). The location of the electrodes were co-registered on The forward model was then computed using the symmetric Boundary Element Method developed in the open OpenMEEG freeware, using default values for conductivity and layer thickness (Gramfort et al., 2010). The covariance matrix computed on a 30 seconds resting state window recorded as first acquisition of the session has been used for each session as noise covariance matrix. For each of the single pulse TMS selected epochs, the source level activation has been computed using a Minimum norm imaging linear method with sLORETA as inverse model. The dipole orientation of the source model has been defined as unconstrained to the cortex surface. Sources orientation was kept orthogonally to the cortical surface and sources amplitude was estimated using the default values of the Brainstorm implementation of the whitened and depth-weighted linear L2-minimum norm solution.

In order to extract local and remote source activity (L/RSA) power, two ROIs were created on each individual anatomy using the individual TMS coordinates for V1 and MT (see above, section “Single pulse TMS”), covering about 195 vertices of cortical mesh for V1 and 320 for MT. LSA and RSA (i.e. power was then computed for each cortical target by averaging the absolute, smoothed (using a spatial smoothing filter with full width at half maximum of 5 mm) and normalized (z-score against baseline) source activity within its corresponding ROI. LSA consisted in the matched condition where local source activity was extracted. RSA consisted in the same procedure but extracting source activity of the non-stimulated area (unmatched condition, see Figure 1D for an illustration). Grand average L/RSA power was finally calculated for each stimulation site and ccPAS condition by averaging L/RSA power across subjects (see Figure 1D for an illustration of the L/RSA concept).

### TMS-EEG Connectivity Analysis

We explored effective connectivity at different frequency ranges using spectral Granger Causality (GC), which is a metric of directed interareal influence (Friston et al., 2014) in a broader visual network known to be involved in motion direction discrimination (Pascual-Leone et al., 2001). Local source activity was extracted from these two additional areas: the inferior parietal sulcus (IPS, ∼120 vertices) and the Frontal Eye Field (FEF, ∼140 vertices).

Granger causality is a measure of linear dependence, which tests whether the variance of error for a linear autoregressive model estimation (AR model) of a signal *x*(*t*) can be reduced when adding a linear model estimation of a second signal *y*(*t*). If this is true, signal *y*(*t*) has a Granger causal effect on the first signal, *x*(*t*) i.e., independent information of the past of *y*(*t*) improves the prediction of *x*(*t*) above and beyond the information contained in the past of *x*(*t*) alone. The term independent is emphasized because it creates some interesting properties for GC, such as that it is invariant under rescaling of the signals, as well as the addition of a multiple of *x*(*t*) to *y*(*t*). The measure of Granger Causality is nonnegative, and zero when there is no Granger causality. according to the original formulation of GC, the measure of GC from *y*(*t*)to *x*(*t*) is defined as:

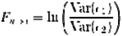

Which is 0 for Var (*e*_1_) = Var (*e*_2_) and a non-negative value for Var (*e*_1_) > Var (*e*_2_). Note Var (*e*_1_) = ≥ Var (*e*_2_) always holds, as the model can only improve when adding new information. always holds, as the model can only improve when adding new information.

Under fairly general conditions, *F* _*y*→ *x*_ can be decomposed by frequency. If the two AR models in time domain are specified as:

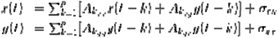

In each equation the reduced model can be defined when each signal is an AR model of only its own past, with error terms and. Then we can defined the variance-covariance matrix of the whole system as:

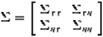

where ∑_*xx*_ = var(*σ*_*xx*_), etc. Applying the the Fourier transform to these equations, they can be expressed as:

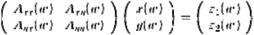

Rewriting this as

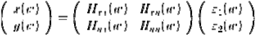

Where *H*(*w*) is the transfer matrix. The spectral matrix is then defined as:

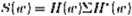

Finally, assuming independence of the signals *x* and *y*, and ∑_*xy*_ *=*∑_*yx*_ *=* 0, we can define the spectral GC as:

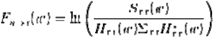

GC was then computed between the four clusters (V1, MT, IPS and FEF), in a frequency range of 1-45 Hz and with a definition of 3 Hz, in order to explore the role of different rhythms in the causal connectivity between brain regions.

Note that the use of Granger causality on source-projected EEG data reduces the problem of signal mixing and volume conduction, probably because Granger causality reflects causal, i.e. time-delayed, interactions, and explicitly discards instantaneous interactions resulting from signal mixing (Bastos et al., 2015; West et al., 2020; Michalareas et al., 2016).

Importantly, we computed GC on the EEG signals in response to single pulse TMS. This combination allowed us to distinguish direct output signals from re-entrant signals (see Figure 1C/D for an illustration). In the manuscript, we used the term “*output connectivity*” to refer to the condition where single pulse TMS was applied to the first area (e.g., TMS over V1, GC from V1 to MT). In contrast, we refer to “*re-entrant connectivity*” for the condition when TMS was applied to the second area (e.g., TMS over MT and GC from V1 to MT).

### Statistics

#### Behavioral data

We individually extracted for the pre and post measurements, the direction range thresholds using all trials (i.e., 150 trials) by fitting a Weibull function, which defined the direction range level at which performance reached 75% correct. These direction range thresholds were then normalized to the maximum possible range of motion (360°), resulting in a normalized direction range threshold (NDR), a procedure previously described (e.g., Huxlin et al., 2009) using the following formula:

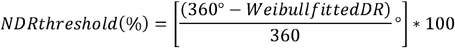

NDR ratios were entered into a mixed ANOVA that included Time (Pre vs Post) and ccPAS type (FW vs BW) as within-subject factor and ccPAS order (first FW vs first BW) as between-subject factor. Post hoc t-tests were performed when appropriate and significance was defined for p values inferior than 5%.

#### EEG data

Significant differences in L/RSA curves as well as in connectivity strength/frequency resolved GC were evaluated within-subjects through a non-parametric, cluster-based corrected, permutation testing (Lage-Castellanos et al., 2010).

#### Behavior and EEG

Individual NDR values were then computed as a ratio expressing the change between Pre and Post and entered into forward stepwise regressions to determine, which neural pathway modulation(s) (V1-to-MT or MT-to-V1) and which frequencies best predicted changes in motion direction discrimination.

## Results

Overall, the 16 participants tolerated well the experiment. The two sessions were equally rated in terms of discomfort and sensations. One participant dropped out, resulting in 16 datasets for FW ccPAS and 15 datasets for BW ccPAS.

### Local and Remote Source activity (LSA & RSA)

Figure 2 shows the results related to the LSA and RSA analyses. LSA refers to the matched condition (source activity extracted locally) and RSA refers to the unmatched condition (source activity extracted from the other region) (see the Methods section for more details). The focus of the study was on LSA and RSA early components (<200ms after the onset).

**Fig. 2:**
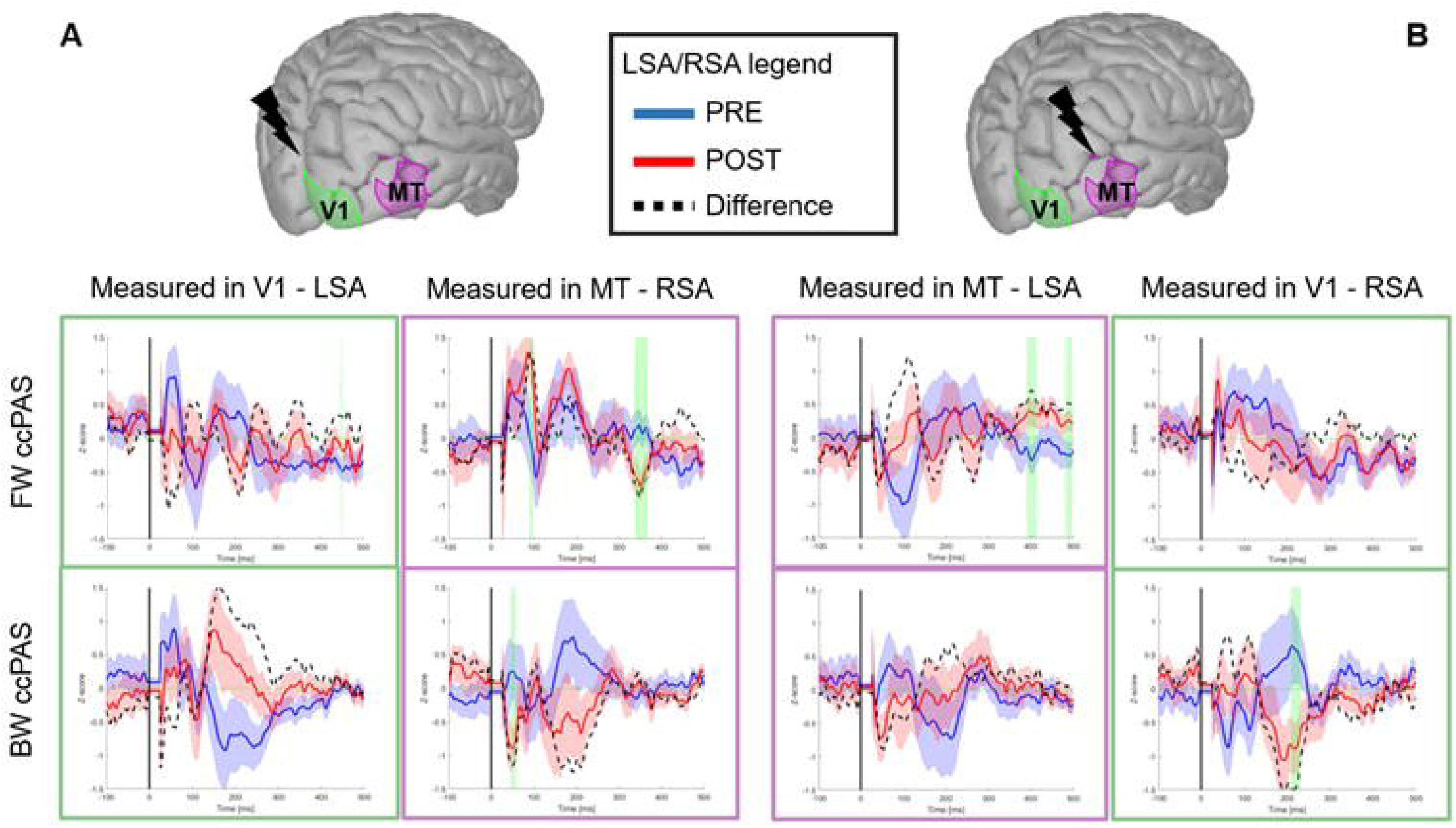
Local and Remote Source Activity on V1 and MT. Clusters of areas V1 and MT are represented on a template brain anatomy. Left (**A**): single pulse TMS over V1. Right (**B**): single pulse TMS over MT. Top row: FW ccPAS condition, Bottom row: BW ccPAS condition. The blue and red shaded area in the plots represent the standard error, while the green shaded areas represent significant difference between POST and PRE recording according to permutation test.

Considering FW ccPAS, LSA over V1 showed a non-significant decrease in local activity (from 15ms to 100ms) and an increase in local MT activity (from 30ms to 150ms). Remote activity remained stable for both V1 and MT.

BW ccPAS LSA profiles over V1 also showed a non-significant decrease in the early components (from 15ms to 100ms), but also a decrease in local MT activity. However, RSA showed a significant decrease between 15ms to 30ms over MT but an increase in the top-down direction, when stimulation was given to MT and EEG was measured in V1 in the later components (from 220ms to 250ms) in line with signal propagation from MT. Of note, Figure 2 also shows an increase of endogenous alpha oscillations in both visual areas triggered by the stimulation.

### Connectivity analysis

Figure 3 reports changes in sGC within the selected areas of the visual network. Spectral representations are only reported for the main pathway of interest (V1-MT).

**Fig. 3:**
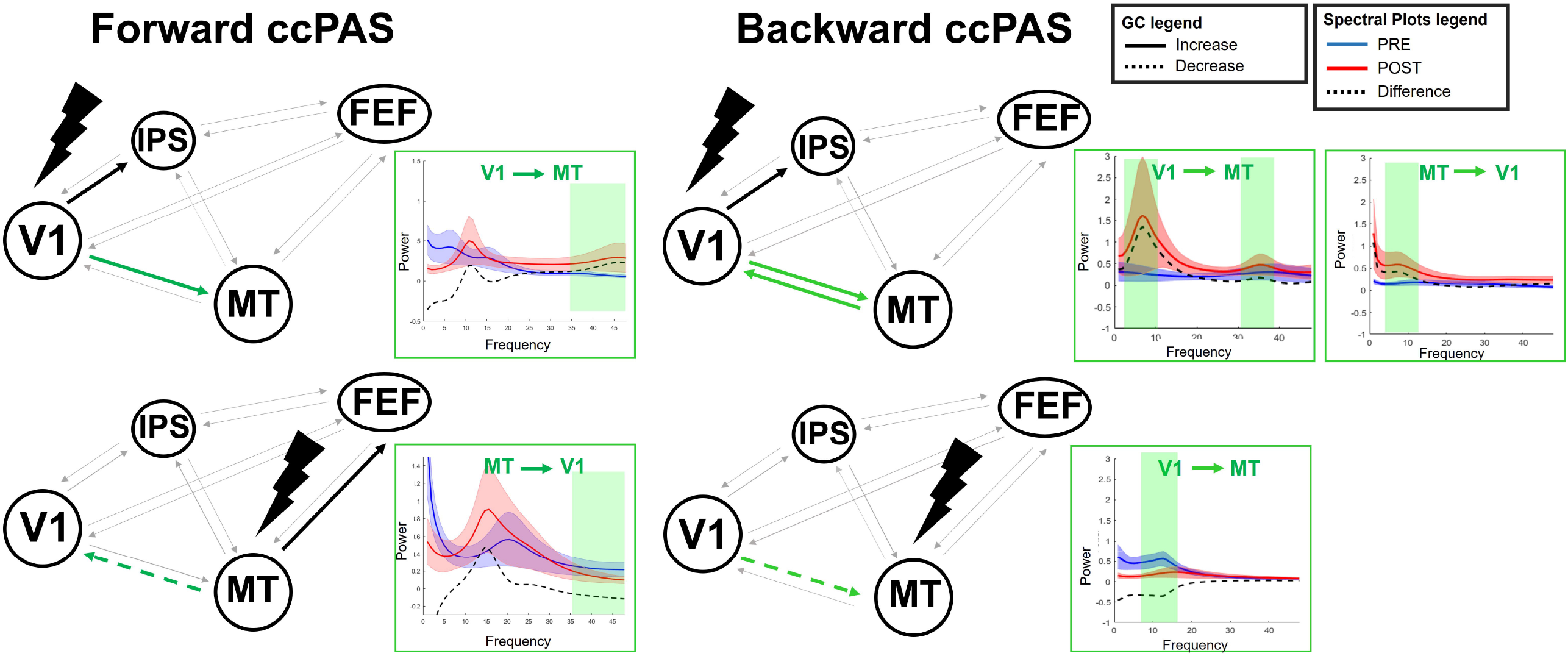
Spectral Granger Causality of the visual network after single pulse TMS on V1 (top row) and MT (bottom row) for FW ccPAS condition (left column) and BW condition (right column) Areas considered for the computation of spectral Granger Causality: V1, MT, FEF and IPS. Spectral plots showing significant changes in GC in the frequency domain are displayed for the V1-MT pathway (other significant connectivity PSD plots are provided in Supplementary Fig. S2). The blue and red shaded areas in the represent the standard error, while the grey shaded areas represent statistically significant differences between PRE and POST GC according to permutation statistical test.

FW ccPAS (Fig.3, left column) showed a significant increase in the bottom-up V1-to-IPS and V1-to-MT connectivity in the gamma band (35-45 Hz), when V1 was stimulated. When MT was stimulated, direct output pathways were modulated: the bottom-up MT-to-FEF connections got significantly promoted in the alpha band (8-12 Hz), while the top-down MT-to-V1 inputs were significantly inhibited in the gamma band (see Supplementary Figure S2).

BW ccPAS (Fig.3, right column), also significantly increased direct bottom-up inputs (V1-to-MT and V1-to-IPS) in the alpha and gamma band when V1 was stimulated. Interestingly the re-entrant MT-to-V1 pathway was also increased in the alpha range (MT-to-V1). When MT was stimulated, the direct top-down inputs to V1 decreased, but the re-entrant V1-to-MT inputs increased, both in the alpha band.

### Behavioural results

We then explored how these changes in local EEG activity and in inter-areal connectivity translated into behavioural changes after the two ccPAS interventions. The normalized direction ratio (NDR) are represented in Figure 4A. The mixed ANOVA on the NDR values showed a main effect of Time (F(1,11) = 13.3, p = 0.004) and a significant ccPAS type by Time (F(1,11) = 5.37, p = 0.04). The post hoc comparisons showed that the pre-post difference was only significant in the BW ccPAS (BW ccPAS: t(13) = 3.95, p = 0.002, FW ccPAS: t(14) = 1.43, p = 0.17, paired t tests). The order effect was not significant (F(1,11) = 1.21, p = 0.29).

**Fig. 4:**
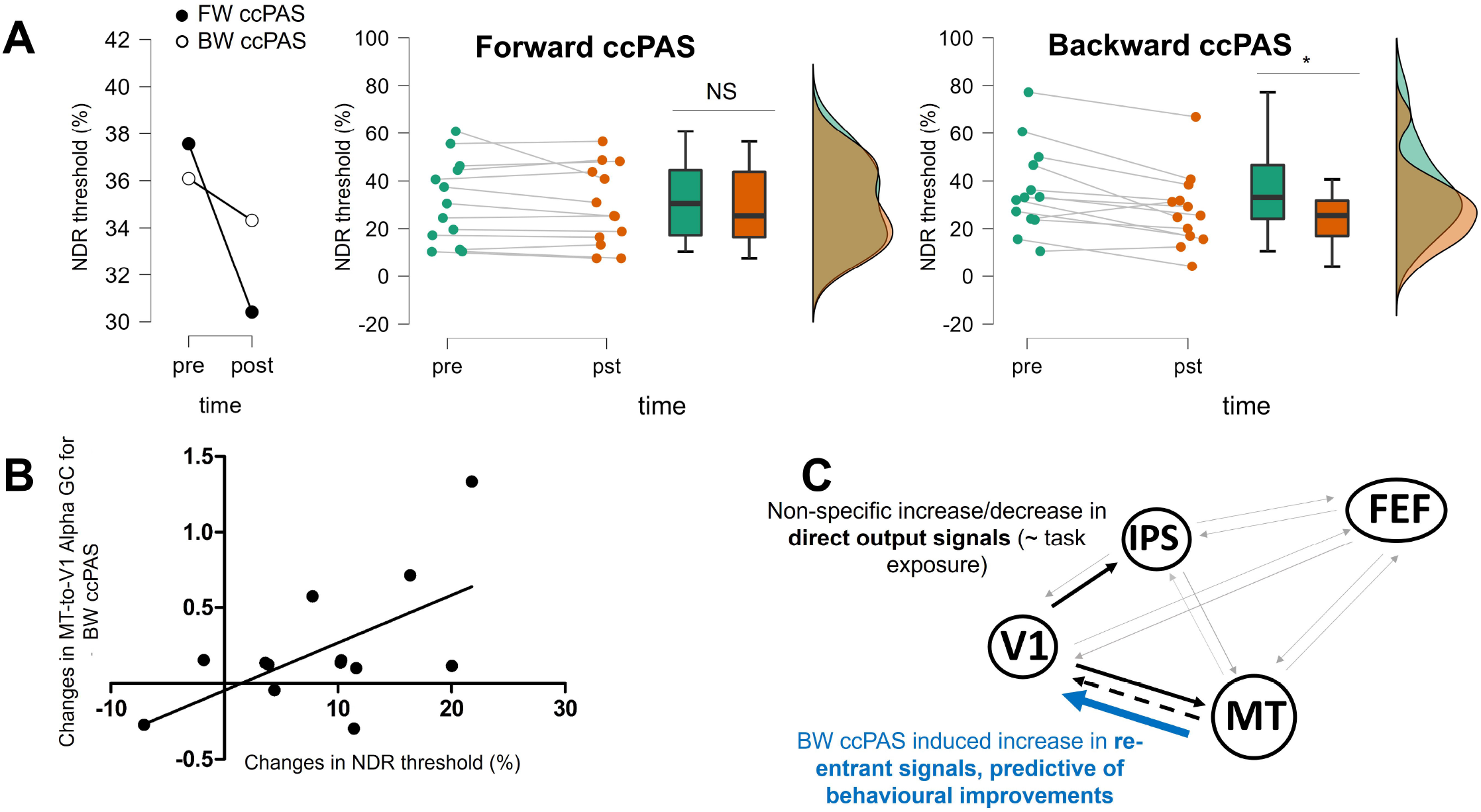
A Group-level changes in NDR threshold (Left panel), individual data and post hoc within group comparison for FW ccPAS (middle panel) and for BW ccPAS (right panel); **B**: Correlational plot between the changes in MT-V1 connectivity strength and the changes in NDR threshold for the BW ccPAS illustrating the results of the forward stepward regression model; **C:** Summary of the connectivity and behavioural results.

Moreover, we compared confidence ratings and here as well, we found a significant Time by ccPAS type interaction (F(1,14) = 4390, p < 0.001), reflecting the specific increase in ratings for the BW ccPAS group (BW ccPAS: t(14) = 1.6, p = 783, FW ccPAS: t(15) = 22, p < 0.001, post hoc within group comparisons).

In order to relate the EEG changes described above to these differences in motion direction discrimination after the two interventions, we designed two forward stepwise regression models, the first one with the baseline-corrected NDR values belonging to the FW ccPAS session and the second was with the baseline-corrected NDR values belonging to the BW ccPAS session as dependent variables. The results showed that the first model (FW ccPAS) was not significant, suggesting that none of the connectivity changes could explain changes in NDR in this ccPAS condition. For the second model (BW ccPAS), the model was significant and the retained variable was the re-entrant MT-to-V1 pathway (see Table 1 and Figure 4B for an illustration of the relationship between performances and MT-to-V1 connectivity changes). These results showed that the significant improvement found after BW ccPAS could be explained by the increase in top-down inputs from MT-to-V1.

**Table 1.**
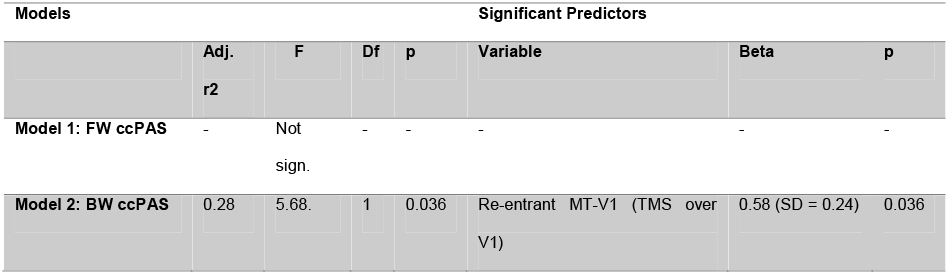
Multiple regression analyses

## Discussion

In the present study, we used TMS-EEG coupling to investigate whether cortico-cortical PAS between V1 and MT can modulate direction specific network plasticity and improve motion direction discrimination in young healthy participants.

### 1 Strengthening MT-to-V1 inputs improves motion direction discrimination and awareness

The intervention of interest tested in this study aimed at strengthening synaptic plasticity of the MT to V1 back-projections. V1 receives inputs from neighbouring area V2 and from a set of higher cortical areas (including MT), transmitting the outcomes of many cognitive operations such as attention, expectation or imagination. As a matter of fact, V1 receives considerably more backward and lateral inputs than forward thalamic afferents (Budd, 1998). V1 is therefore a processing and integrative centre of a complex cortical processing cascade. Those modulatory backward projections from higher visual areas or associative areas control the gain of thalamocortical inputs to V1 through the activation of glutamatergic receptors (Hupé et al., 1998; Ekstrom et al., 2008; Muckli et al., 2013).

Evidence from macaques (Lamme et al. 1998) and humans (Pascual-Leone et al. 2001; Silvanto et al. 2005) has shown that back projections from extrastriate areas to the primary visual area (V1) are crucial for motion discrimination. In an earlier study, Romei and colleagues (2016) applied MT-V1 ccPAS (BW ccPAS) on healthy subjects and found improved motion coherence discrimination, suggesting plasticity changes within this pathway. We provided further evidence for this finding with a significant enhancement of motion direction discrimination only after BW ccPAS. Furthermore, BW ccPAS significantly improved motion awareness, as evidenced by the increased metacognitive confidence rating, confirming the role of this pathway for motion awareness (Pascual-Leone et al. 2001; Silvanto et al. 2005). Of note, confidence rating has been shown to influence visual discrimination (Bonder et al., 2019). Therefore, we cannot infer whether ccPAS exerted its effects directly on motion perception, which then impacted on the confidence rating or whether it is the opposite. Interestingly, previous research has found that the macaque lateral intraparietal area (LIP), homologue of the human intraparietal sulcus (IPS) plays a crucial role in forming confidence in perceptual decision-making (Huk et al. 2017; Pasternak et al., 2020). Granger Causality results did reveal an increase in V1-to-IPS connectivity for both conditions, as well as source activity in the IPS (see supplementary Figure S2). Therefore, strengthening the MT-to-V1 pathway might also support the neural circuitry of metacognitive judgements of perceptual decision-making, but future studies should be performed to understand how these two brain functions interact with each other.

In contrast, modulating the reciprocal forward projections did not further boost motion direction discrimination or motion awareness. The FW ccPAS only showed a small, not significant improvement, that might be related to a well-reported gradual improvement through practice at motion direction discrimination (Gibson, 1963; Sagi, 2011). Importantly, the absence of significant improvements justifies the absence of a Sham comparison in this study. This lack of improvements in motion discrimination for the FW ccPAS condition is partly supported by anatomical studies, showing a mixing of early parallel pathways within V1 that result in an apparent lack of compartmentalization in outputs from V1 to extrastriate cortical areas (Sincich et al., 2002, Xiao et al., 2004). Some researchers have even provided evidence against parallel-processing models of the primate visual system. Furthermore, the fact that the ccPAS protocol was applied at rest might prevent the possibility of a functional routing through the specific cortical circuits relevant to motion processing. Finally, neurons projecting directly from V1 to MT are located within layer 4B, closer to the layer 4Cα border, deeper than neurons projecting to other visual areas such as V2 or V3 (Nassi et al., 2007) and deeper than MT-projecting neurons to V1, mostly located within layer 1 (Blasdel and Lund, 1983). This could explain why the forward inputs are less sensitive to TMS.

### 2 The neural correlates of ccPAS

Congruent with the concept of network-based interventions, none of the ccPAS condition resulted in local changes in EEG activity (denoted LSA in the present manuscript). However, when EEG responses to TMS were measured in the opposite region (i.e., TMS over V1, EEG response measured in MT, labelled RSA in the paper) we found significant changes that we interpreted as reflecting changes in signal propagation from one region to another one. FW ccPAS did not show any significant difference, but we found a decrease in top-down RSA (MT to V1) for the BW ccPAS protocol. The GC results provided additional insights on synaptic transmission and information flow within the cortical motion network. While the L/RSA profiles are computed from a mixture of signals from all frequency bands, we used spectrally resolved GC (Chicharro, 2012) to test hypotheses on specific oscillatory channels mediating backward and forward inputs (van Kerkoerle et al., 2014; Bastos et al., 2015). Crucially, the combination of GC measures with single pulse TMS allowed us to distinguish between direct output signal diffusion from re-entering signal transmission (Winkler et al., 2015). This distinction was particularly relevant to the BW ccPAS condition were the re-entrant top-down MT-to-V1 connection was significantly increased in the Alpha range (8-12Hz), in proportion with enhanced motion direction discrimination (see Figure 4C for a summary of our findings). In turn, the bottom-up, re-entrant V1-to-MT pathway was decreased in Alpha, suggesting that the BW ccPAS protocol is highly sensitive to and primarily acts on re-entrant fibers (Fig.3, right column). These findings are in accordance with animal and human electrophysiological studies: Alpha oscillations in the visual cortex have been shown to characterize backward processing while gamma waves are thought to mediate forward connectivity direction studies (van Kerkoerle et al., 2014; Michalareas et al., 2016, Richter et al., 2018). The changes in inter-areal coupling we report in the present study might support the hypothesis of a better local and specialized processing after the BW ccPAS protocol.

### 3 Non-specific changes potentially attributable to task exposure

Local EEG source activity in response to TMS over V1 showed a common, non-significant decrease in early components (from 20ms to 80ms) while sGC revealed that both ccPAS interventions induced an increase in direct output connectivity from V1-to-MT and from V1-to-IPS, all in the Gamma band except V1-to-IPS for the FW ccPAS condition which occurred in the Alpha band. These increases were not predictive of any behavioural improvement. Studies (e.g., Koch et al., 2007) showed that during top-down attention the synchrony between neurons in prefrontal and parietal areas is stronger in lower frequencies; whereas a stronger coherence at higher frequency bands is observed during bottom-up attention. This might indicate that bottom-up, exogenous attention increases with task exposure, possibly mediated by an increase in Gamma oscillations. These non-specific increases fit with the visually triggered increase of information flow with task exposure, specific to low level visual perception that has been associated with purely externally driven visual experience (Busch et al., 2006, Zahoping, 2008, Katsuki et al., 2013; Dijkstra et al., 2017; Regev et al., 2018; Jia et al., 2020).

### 4 Perspectives

There is evidence that the effect and the magnitude of spike-timing dependent plasticity (STDP) generated by the ccPAS intervention over the MT-to-V1 back projections might be subject to state-dependent shifts in neocortical excitability (Chao et al. 2015; Veniero et al. 2013; Fehér et al. 2017). The different excitability states could be indexed or indirectly read out using EEG-derived phases of neocortical oscillations, in particular during the Alpha cycle (Fehèr et al., 2017). Therefore, to further improve the effects of the BW ccPAS intervention, one could implement it in a neocortical excitability state-dependent framework. Future studies should test whether phase information extracted through online EEG recordings could be used to trigger the paired pulse TMS over MT and V1, either to control the onset of the paired pulse or the time delay between the conditioning and test pulse (MT and V1 pulse). If ccPAS effects depend on the in-going cortical excitability state, the size and effects of ccPAS might be phase dependent.

Finally, BW ccPAS effects in the visual system could be used to quantify the acute capacity of the visual system to reorganize and from this index, extract a predictor of visual recovery after a visual stroke for instance. The “amount” of induced plasticity in hemianopic stroke patients could reveal precious information about the functional state of their visual system, to predict whether an individual brain is able to recruit backward projecting neurons to the spared V1 population to support recovery.

## Abbreviations

BOLD: Blood oxygenation level dependent
ccPAS: Cortico-cortical paired associative stimulation
EEG: Electroencephalography
FEF: Frontal eye fields
fMRI: Functional magnetic resonance imaging
IPS: Intraparietal sulcus
LTP: Long term potentiation
TMS: Transcranial magnetic stimulation
V1: Primary visual cortex
MT: Medio-temporal cortex

---

**Fig. S1:**
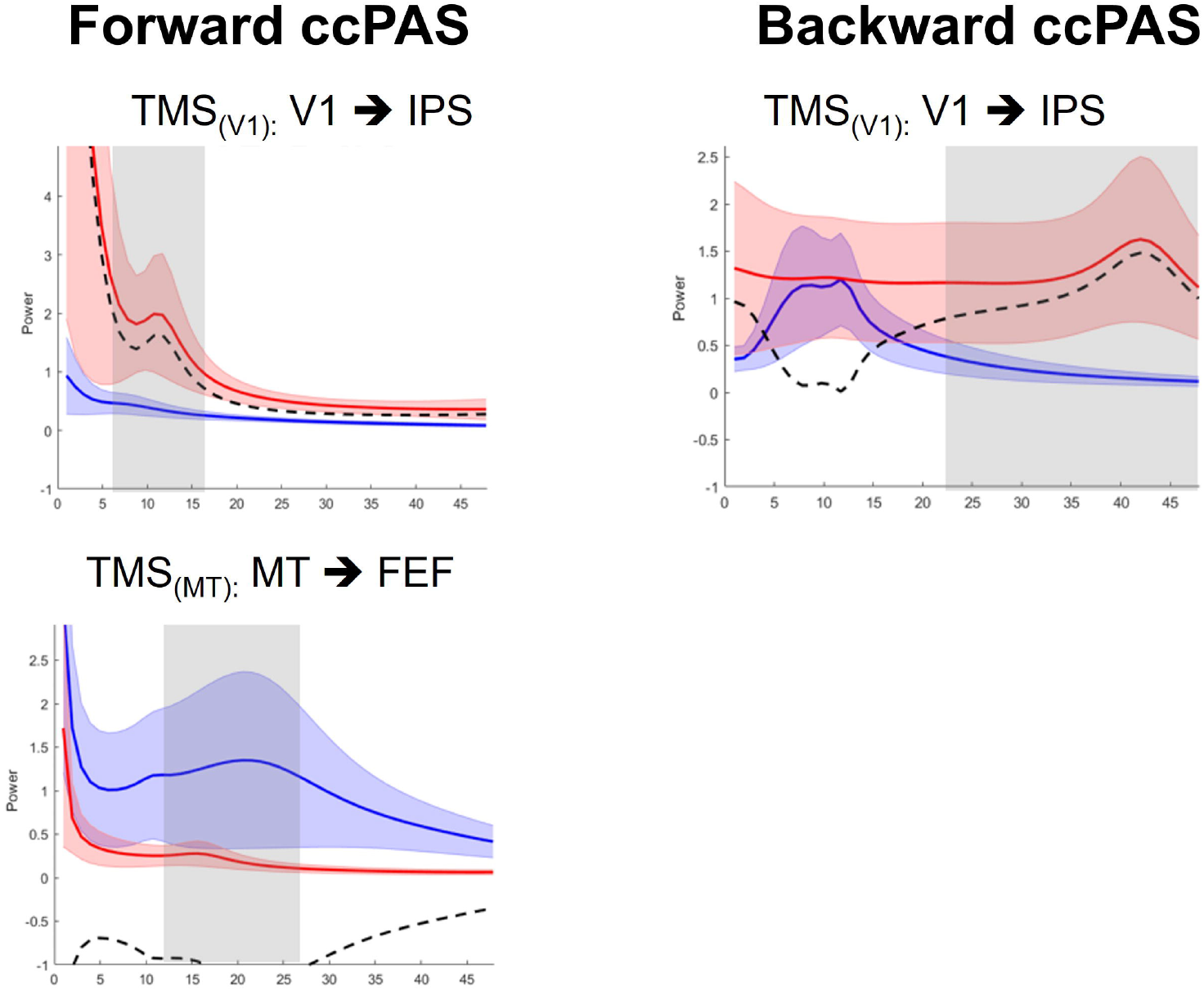
Additional significant connectivity changes measured with spectrally resolved Granger Causality after single pulse TMS for FW ccPAS (left column) and BW ccPAS (right column).

**Figure S2:**
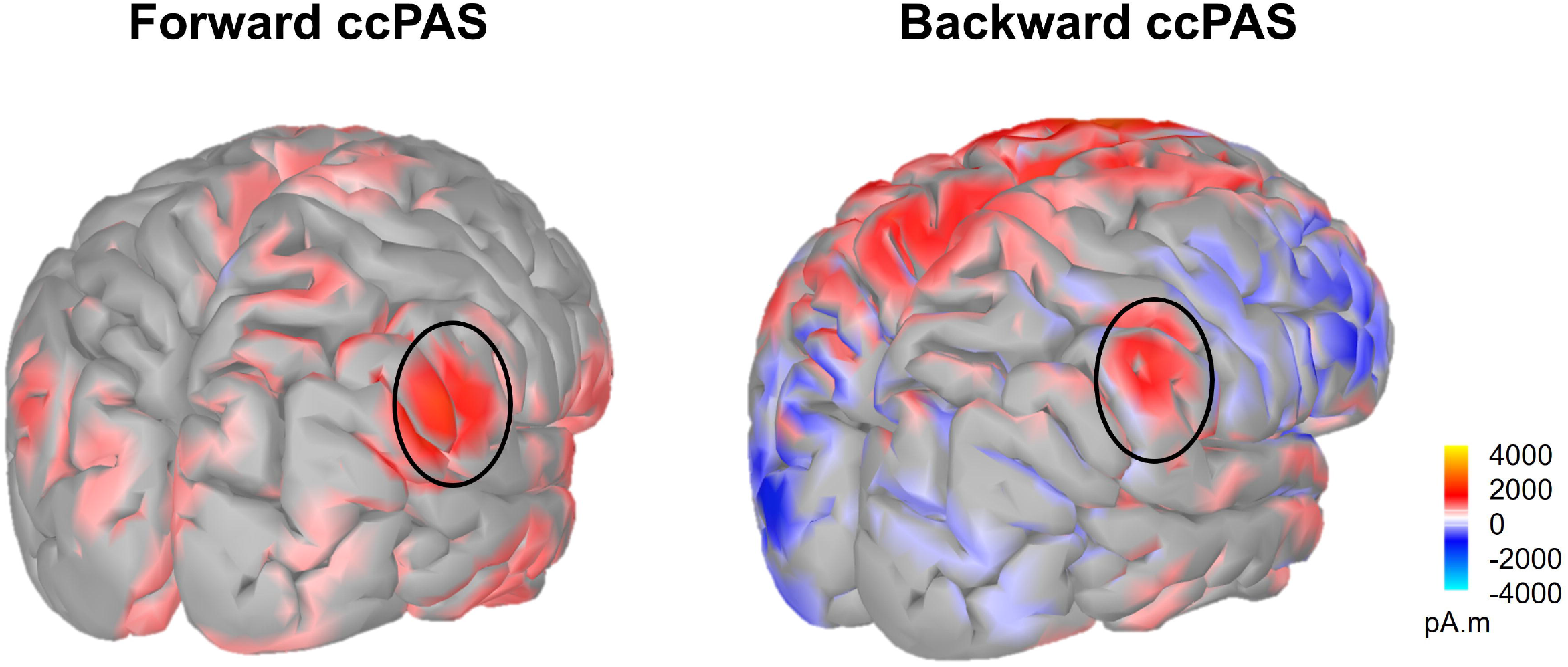
Source activity of the contrast post ccPAS > pre ccPAS showing the increased activity in the IPS (inferior parietal sulcus) for the FW ccPAS condition (right) and for the BW ccPAS condition (left) after single pulse TMS.

